# Galectin-8 senses phagosomal damage and recruits selective autophagy adapter TAX1BP1 to control *Mycobacterium tuberculosis* infection in macrophages

**DOI:** 10.1101/2020.06.30.180877

**Authors:** Samantha L. Bell, Kayla L. Lopez, Jeffery S. Cox, Kristin L. Patrick, Robert O. Watson

## Abstract

*Mycobacterium tuberculosis* (Mtb) infects a quarter of the world and causes the deadliest infectious disease worldwide. Upon infection, Mtb is phagocytosed by macrophages and uses its virulence-associated ESX-1 secretion system to modulate the host cell and establish a replicative niche. We have previously shown the ESX-1 secretion system permeabilizes the Mtb-containing phagosome and that a population (~30%) of intracellular Mtb are recognized within the cytosol, tagged with ubiquitin, and targeted to the selective autophagy pathway. Despite the importance of selective autophagy in controlling infection, the mechanisms through which macrophages sense and respond to damaged Mtb-containing phagosomes remains unclear. Here, we demonstrate that several cytosolic glycan-binding proteins, known as galectins, recognize Mtb-containing phagosomes. We found that galectins-3, -8, and -9 are all recruited to the same Mtb population that colocalizes with selective autophagy markers like ubiquitin, p62, and LC3, which indicates Mtb damages its phagosomal membrane such that cytosolic host sensors can recognize danger signals in the lumen. To determine which galectins are required for controlling Mtb replication in macrophages, we generated CRISPR/Cas9 knockout macrophages lacking individual or multiple galectins and found that galectin-8^-/-^ and galectin-3/8/9^-/-^ knockout macrophages were similarly defective in targeting Mtb to selective autophagy and controlling replication, suggesting galectin-8 plays a privileged role in anti-Mtb autophagy. In investigating this specificity, we identified a novel and specific interaction between galectin-8 and TAX1BP1, one of several autophagy adaptors that bridges cargo and LC3 during the course of autophagosome formation, and this galectin-8/TAX1BP1 interaction was necessary to efficiently target Mtb to selective autophagy. Remarkably, overexpressing individual galectins increased targeting of Mtb to antibacterial autophagy and limited Mtb replication. Taken together, these data imply that galectins recognize damaged Mtb-containing phagosomes, recruit downstream autophagy machinery, and may represent promising targets for host-directed therapeutics to treat Mtb.

## INTRODUCTION

*Mycobacterium tuberculosis* (Mtb), which causes tuberculosis, infects approximately 10 million people annually and kills about 1.5 million, making it the deadliest infectious disease worldwide (World Health Organization, 2019). Spread in aerosolized droplets when an infected person coughs, Mtb travels to the depths of the lungs where it is phagocytosed by alveolar macrophages. Typically, macrophages are incredibly efficient at identifying and destroying invading microbes, and they have numerous potent killing mechanisms, including lysosomal degradation, reactive oxygen species, antimicrobial peptides, guanylate-binding proteins (GBPs), and autophagy (Weiss & Schaible, 2015). However, Mtb employs strategies to resist nearly all of these defense mechanisms and survives and replicates in macrophages (Kaufmann & Dorhoi, 2016; Upadhyay et al., 2018). Understanding the few mechanisms by which macrophages can successfully control Mtb is critical for the future development of effective therapies for this difficult-to-treat pathogen.

One way a macrophage can control intracellular Mtb is through selective autophagy, a specific form of autophagy whereby a cell tags unwanted cytosolic cargo with ubiquitin, which serves as an “eat me” signal (Boyle & Randow, 2013; Khaminets et al., 2016; Stolz et al., 2014). Ubiquitin-tagged cargo can then be coated by a variety of selective autophagy adapters (p62/SQSTM1, Calcoco2/NDP52, Optineurin/OPTN, etc.), which have ubiquitin-binding domains that promote their recruitment to tagged cargo. These adapters also have an LC3 interaction region (LIR), a motif that enables binding to the autophagy protein LC3 and the closely related GABARAP proteins (Wild et al., 2014). As a result, selective autophagy adapters serve as bridges between ubiquitinated cargo and the LC3-decorated autophagophore that will ultimately engulf and degrade the cargo. Numerous types of cargo, including damaged mitochondria (mitophagy), protein aggregates (aggrephagy), and cytosolic pathogens (xenophagy), can be degraded via selective autophagy, and various subsets of adapters are associated with different types of cargo. For example, mitophagy utilizes NDP52 and OPTN, aggrephagy p62 and NBR1, and xenophagy p62 and NDP52 (Farré & Subramani, 2016; Stolz et al., 2014). However, the biology underlying the redundancy and specificity of these adapters remains poorly understood.

Several lines of evidence indicate that selective autophagy is required for controlling Mtb infection. Our work and that of others have shown that ESX-1-dependent permeabilization of the Mtb phagosome allows the cytosolic DNA sensor cGAS to detect bacterial dsDNA, which triggers both a pro-bacterial type I interferon (IFN) transcriptional response (via the STING/TBK1/IRF3 signaling axis) and anti-bacterial selective autophagy (Collins et al., 2015; Manzanillo et al., 2012; Wassermann et al., 2015; Watson et al., 2012, 2015). Specifically, within 4-6 h after infection, approximately 30% of intracellular Mtb bacilli are surrounded by ubiquitin, LC3, and several selective autophagy adapters (Watson et al., 2012). In the absence of selective autophagy targeting (i.e., adapter-deficient macrophages), Mtb survives and replicates to a higher degree (Watson et al., 2012). While the precise nature of the ubiquitination of Mtb is unclear, several E3 ligases, including Parkin, Smurf1, and TRIM16 colocalize with a subset of Mtb phagosomes and are required for optimal tagging of Mtb with ubiquitin (Chauhan et al., 2016; Franco et al., 2017; Manzanillo et al., 2013). These E3 ligases are required for controlling Mtb replication in macrophages, and Parkin and Smurf1 are further required for controlling Mtb infection *in vivo* in mouse models of infection. Likewise, macrophages lacking the core autophagy protein ATG5 fail to control Mtb replication, and mice with a macrophage-specific ATG5 deletion are incredibly sensitive to Mtb infection and succumb within weeks (Watson et al., 2012). A subsequent report found that ATG5 plays a critical role in neutrophil-mediated inflammation, suggesting autophagy functions in both cell-intrinsic and cell-extrinsic immune responses (Kimmey et al., 2015).

We are continuing to understand the function, impact, and scope of selective autophagy in controlling Mtb infection, and the precise mechanisms used by macrophages to detect damaged Mtb phagosomes and intracellular Mtb bacilli are remain poorly defined. Our previous studies have found that cytosolic DNA sensing through cGAS/STING/TBK1 is required for recognition and targeting; macrophages lacking cGAS or STING target half as many Mtb bacilli to selective autophagy (Watson et al., 2015). However, because a sizable population of Mtb are targeted even in the absence of DNA sensing, it is likely that additional “danger signals” (e.g., microbes or damage caused by microbes) and “danger sensors” are employed by macrophages during Mtb infection (Vance et al., 2009).

One class of danger sensors are galectins, which are a large, highly conserved family of proteins that bind to glycosylated proteins and lipids via their carbohydrate recognition domains (CDRs) (Rabinovich & Toscano, 2009; van Kooyk & Rabinovich, 2008; Vasta, 2009). Despite having no classical secretion signal, many galectins are extracellular where they can bind to glycosylated proteins and lipids on cell surfaces or in the extracellular matrix to modulate cellular processes like signaling, adherence, and migration (Rabinovich & Toscano, 2009; van Kooyk & Rabinovich, 2008). Several galectins are also found in the cytosol where they exert other functions, including acting as soluble receptors for endosomal or lysosomal membrane damage. After disruption of membranes, galectins can access and bind to glycans within the lumen of damaged membrane-bound compartments (Boyle & Randow, 2013; Khaminets et al., 2016; Thurston et al., 2012). Often, intracellular bacteria inflict this type of endosomal damage, and galectins-3, -8, and -9 have been found to colocalize with several intracellular pathogens, including *Salmonella* Typhimurium, *Shigella flexneri, Listeria monocytogenes, Legionella pneumophilia*, and *Yersinia pseudotuberculosis* (Feeley et al., 2017; Thurston et al., 2012). In some cases, the functional consequences of galectin recruitment to intracellular bacteria have been characterized. During *L. pneumophilia* and *Y. pseudotuberculosis* infection, galectin-3 promotes the recruitment of antibacterial GBPs to bacteria, and during *S*. Typhimurium infection of HeLa cells, galectin-8 recruits NDP52, which brings autophagy machinery to exposed bacteria. While some of these pathways have been studied in detail, the specific molecular mechanisms by which macrophages use galectins to detect and target Mtb to selective autophagy have not been fully characterized.

Here we show that galectins-3, -8, and -9 are recruited to Mtb in macrophages, and that galectin+ bacteria are the same population targeted to selective autophagy. Deletion of galectin-8, but not galectins-3 or -9, decreased targeting of Mtb as monitored by LC3 recruitment and by bacterial survival/replication. Deleting all three galectins did not amplify these phenotypes, suggesting galectin-8 is the most crucial for recognition and targeting of Mtb in macrophages. Using immunoprecipitation and mass spectrometry, we found that galectin-8 interacts with the selective autophagy adapter TAX1BP1, but this interaction is independent of TAX1BP1’s ubiquitin-binding domain. Furthermore, in Mtb-infected macrophages, we found that the recruitment of TAX1BP1 to Mtb required both its interaction with galectin-8 and its ubiquitin-binding domain. Finally, we found that overexpression of galectins-8 and -9 significantly augmented the ability of macrophages to control Mtb survival and replication, indicating that while specific galectins may not be essential for targeting Mtb to selective autophagy, they are sufficient, which raises the possibility of targeting this detection and destruction pathway for the development of future host-directed therapies.

## RESULTS

### Galectins-3, -8, and -9 access the lumen of damaged Mtb-containing phagosomes to detect and target cytosolically exposed bacilli

Because galectins have previously been implicated in sensing phagosomal damage (Feeley et al., 2017; Thurston et al., 2012), we hypothesized that they may play a role in sensing Mtb in macrophages. To test if galectins were recruited to Mtb phagosomes early after infection, we generated 3xFLAG-tagged expression constructs of four different galectins: galectins-1, -3, -8, and -9 (Fig. S1A-B). Galectins-3, -8, and -9 were chosen based on their post-translational modifications during Mtb infection (Budzik et al., 2020; Penn et al., 2018) and because previous studies in non-immune cells have found these galectins colocalized with intracellular pathogens (Thurston et al., 2012). Galectin-1 was chosen as a negative control. We stably expressed epitope-tagged galectins in RAW 264.7 cells, a murine macrophage-like cell line that are a common *ex vivo* infection model for Mtb since they are genetically tractable and respond robustly to Mtb infection (Hoffpauir et al., 2020; Watson et al., 2015). Using these cell lines, we infected with mCherry-expressing Mtb (Erdman strain), and at various times post-infection (3, 6, 12, 24 h), fixed coverslips and used immunofluorescence microscopy to assess 3xFLAG-galectin localization relative to intracellular Mtb (Fig. 1A-B). Galectins-8 and -9, and to a lesser extent galectin-3, were recruited to a sizeable population of Mtb, while galectin-1 was not. Colocalization was detectable at 3 h post-infection and reached a maximum of ~45% galectin-8- or galectin-9-positive bacilli after 24 h. Galectin-3 was recruited to Mtb with similar dynamics, but was only recruited to a maximum of ~20% of bacilli after 24 h. Galectin-1 did not colocalize with Mtb at any time point examined, making it a useful negative control for future experiments.

**Figure 1.**
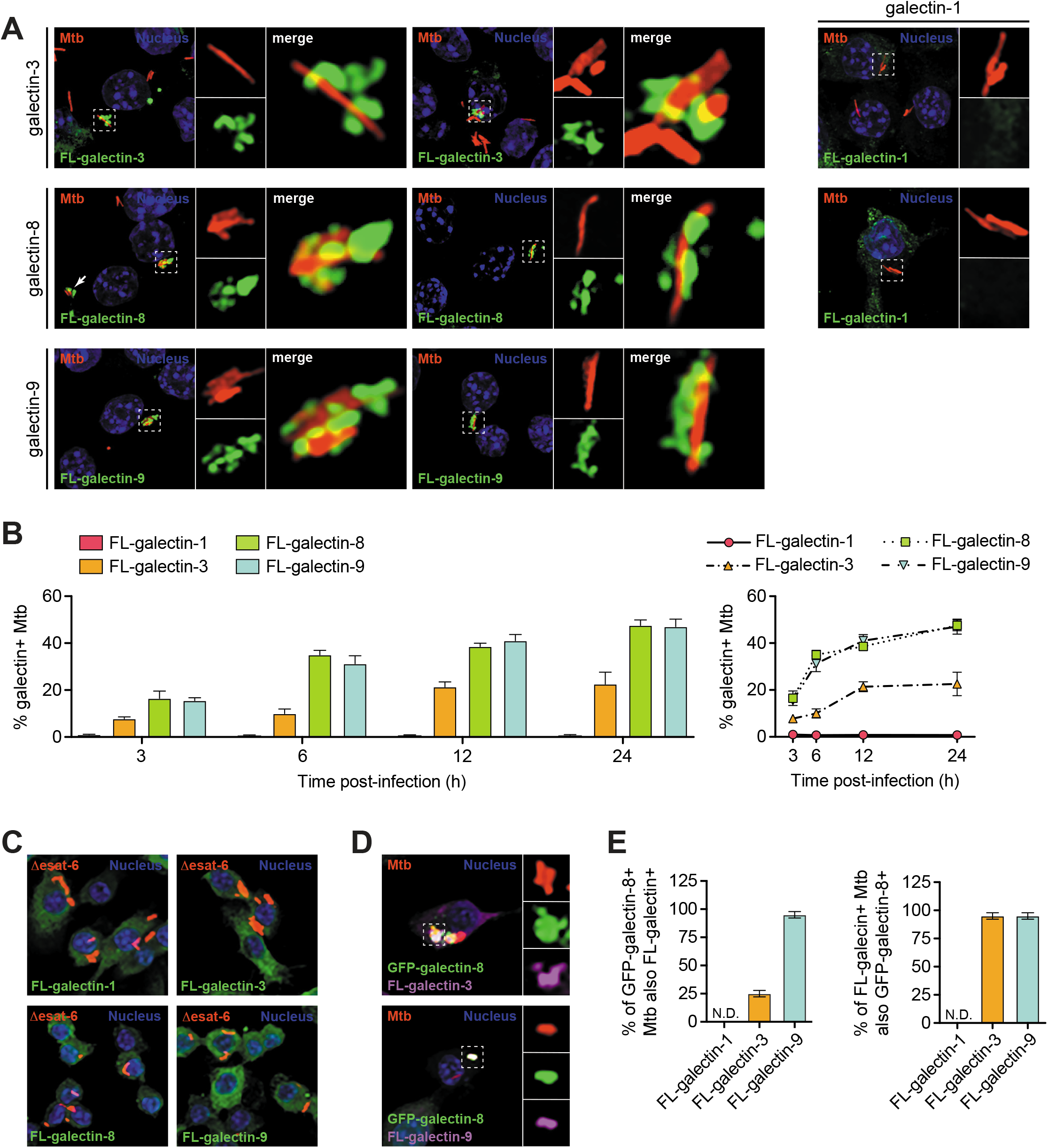
Galectins are recruited to Mtb-containing phagosomes. **(A)** Immunofluorescence of RAW 264.7 cells stably expressing 3xFLAG (FL)-tagged galectins infected with wild-type (WT) mCherry-expressing Mtb (MOI = 1) 6 hr post-infection. Green, FL-galectin; red, mCherry Mtb; blue, DAPI. **(B)** Quantification of FL-galectin-positive Mtb (of indicated genotype) as shown in (A) at indicated timepoints. **(C)** As in (A) but with cells infected with Δesat-6 mCherry-expressing Mtb. **(D)** Immunofluorescence of RAW 264.7 cells stably co-expressing GFP-galectin-8 and FL-galectin-3 or -9 infected with WT mCherry-expressing Mtb (MOI = 1) 6 hr post-infection. Green, GFP-galectin-8; magenta, FL-galectin; red, mCherry Mtb; blue, DAPI. **(E)** Quantification of GFP-galectin-8-positive and FL-galectin-positive Mtb shown in (D). GFP-galectin-8-positive Mtb that are also FL-galectin-3 or -9-positive (left) and FL-galectin-3 or -9-positive Mtb that are also GFP-galectin-8-positive (right). Error bars indicate S.D. of three coverslips per cell line in which at least 100 bacteria were assessed.

Next, we tested if the ESX-1 secretion system, and therefore phagosome permeabilization, was required for galectin recruitment. To do this, we infected 3xFLAG-galectin cells with mCherry-expressing Δesat-6 Mtb (missing a key component for forming pores in the phagosomal membrane (Jonge et al., 2007)). Using immunofluorescence microscopy at 6 h post-infection, we did not observe colocalization of any galectin with Δesat-6 Mtb (Fig. 1C), which indicates that phagosomal permeabilization is required for galectin recruitment. Here and in future experiments, we examined the 6 h post-infection time point since this was the earliest that we observed peak galectin recruitment to Mtb (Fig. 1B). Together, these findings show that ESX-1-induced phagosomal damage is extensive enough to allow cytosolic proteins to access the lumen of the Mtb-containing phagosome.

We next tested whether galectins-3, -8, and -9 were all recruited to the same Mtb-containing phagosomes. To do this, we stably co-expressed GFP-galectin-8 and 3xFLAG-galectin-3 or -9 in RAW 264.7 cells and again infected them with mCherry Mtb. We found that galectin-8 and -9 colocalized in almost all instances (Fig. 1D-E). Likewise, galectin-3 was present on almost all galectin-8+ Mtb, but a large portion of galetin-8+ Mtb did not have galectin-3 present (Fig. 1D-E). This suggests that the same ~30% population of intracellular Mtb accumulates galectins-8 and -9, and sometimes galectin-3.

Based on previous reports and the size of the galectin+ Mtb population, we hypothesized that galectin+ Mtb-containing phagosomes would be positive for selective autophagy markers. To test this, we co-stained for 3xFLAG-glectin-8 and a panel of selective autophagy markers, including ubiquitin (the “eat me” signal), p62 (a selective autophagy adapter), and LC3 (the autophagosome marker). As predicted, the galectin-8+ Mtb were also positive for ubiquitin, p62, and LC3 at 6 h post-infection (Fig. 2). This indicates that galectin+ Mtb are indeed the same population of Mtb that are targeted to selective autophagy.

**Figure 2.**
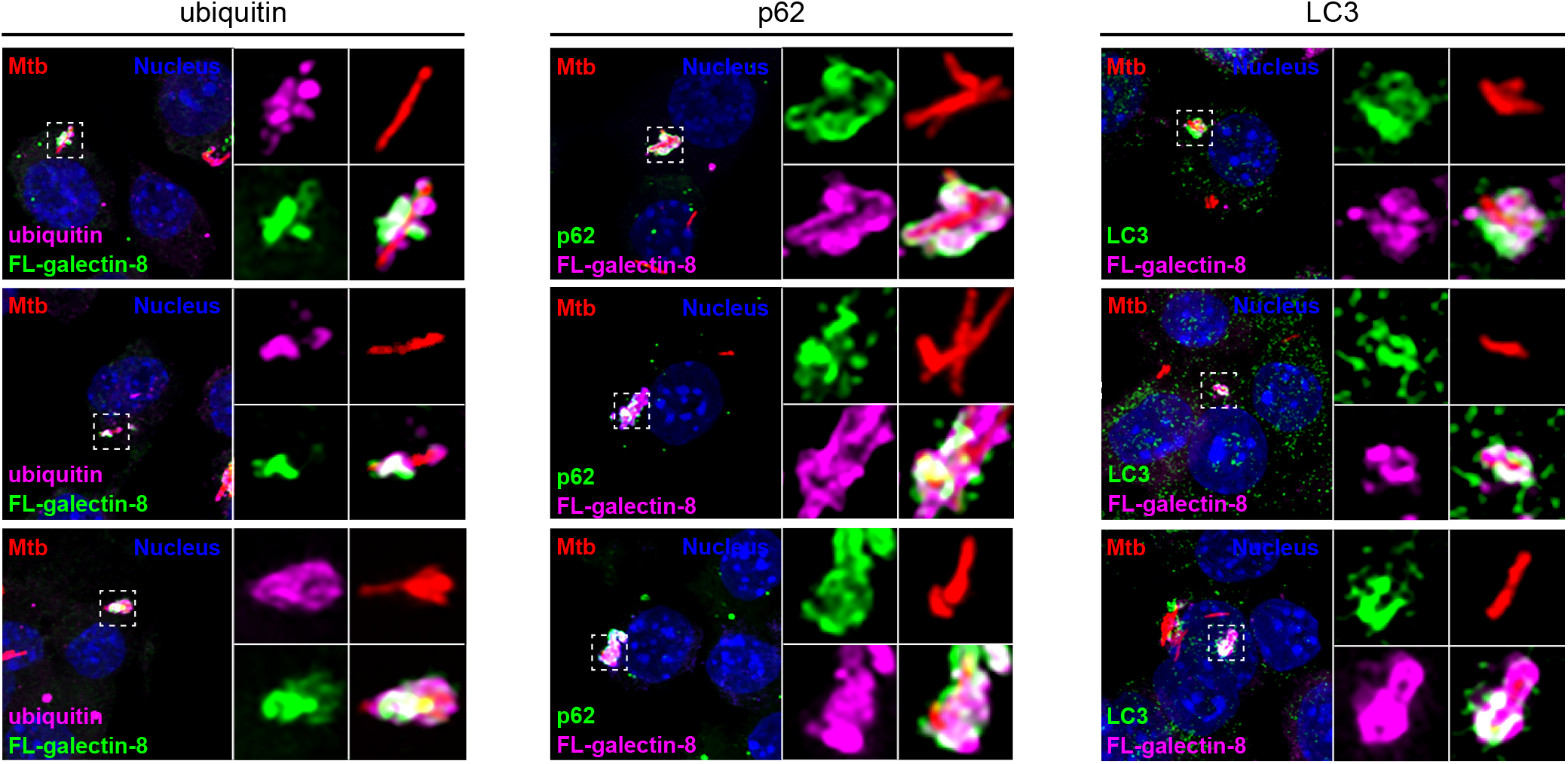
Galectin-decorated Mtb-containing phagosomes colocalize with selective autophagy markers. Immunofluorescence of RAW 264.7 cells stably expressing 3xFLAG (FL)-tagged galectin-8 infected with WT mCherry Mtb (MOI = 1) 6 hr post-infection co-stained for indicated selective autophagy marker (ubiquitin, p62, and LC3). Green and magenta, endogenous selective autophagy marker or FL-galectin-8 (as indicated); red, mCherry Mtb; blue, DAPI.

### Loss of galectin-8 decreases targeting of Mtb to selective autophagy

We next sought to determine if the recruitment of galectins is required for targeting Mtb to antibacterial selective autophagy. To do this, we used a lentiviral CRISPR/Cas9 system to mutate the genes encoding galectins-3, -8, or -9 (*Lgals3, Lgals8, Lgals9*) in RAW 264.7 cells. We designed small guide RNAs (sgRNAs) targeting the first one to two coding exons of each galectin gene; we used GFP-targeted sgRNAs as negative controls. After transducing RAW 264.7 cells stably expressing FLAG-Cas9 with lentiviral sgRNAs constructs, we antibiotic-selected cells, isolated clonal populations, and validated homozygous mutation by sequencing the targeted region. We chose clonal populations that had one or two basepair insertions or deletions that resulted in frameshift mutations early in the transcript (exon 1 or 2) (Fig. 3A). To limit the possibility of off-target and bottleneck effects, we used at least three clonal populations for each gene, and these were derived from two different gRNAs per gene. Since we were unable to identify commercial antibodies that reliably detected the three mouse galectins, we further validated loss of gene expression in the knockout cell lines using RT-qPCR since the mutated transcripts should be degraded via nonsense mediated decay. As expected, all of the knockout cell lines had significantly diminished mRNA expression of the sgRNA-targeted galectin (Fig. S1C).

**Figure 3.**
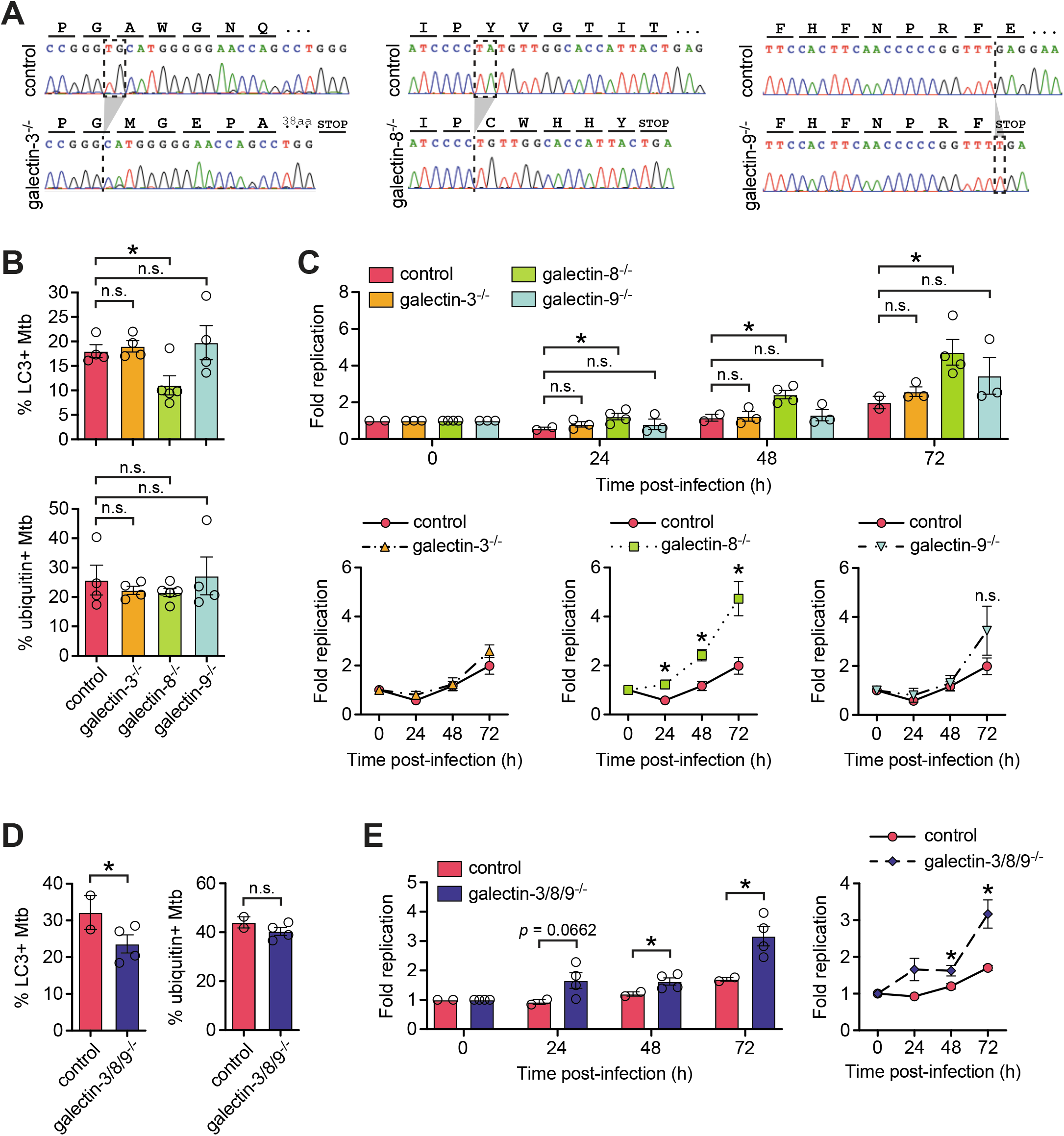
Galectin-8 is required to efficiently target Mtb to selective autophagy to control Mtb replication in macrophages. **(A)** Representative chromatograms from galectin-3, -8, and -9 knockout cells lines indicating the nature of nonsense mutations introduced via CRISPR/Cas9. **(B)** Quantification of LC3-positive (top) and ubiquitin-positive (bottom) Mtb in control (sgRNAs targeting GFP) or individual galectin knockout RAW 264.7 cell lines at 6 h post-infection. Circles represent data for each clonally selected cell line (at least two cell lines per sgRNA and two sgRNAs per galectin gene). **(C)** Fold replication of luxBCADE Mtb (MOI = 1) in control and galectin knockout cell lines at indicated time points. Data normalized to t=0 h. **(D-E)** As in (B-C) but with RAW 264.7 cell lines in which all three galectins are knocked out. Error bars indicate S.E.M. of knockout cell lines, for IF, at least 300 bacteria per cell line were assessed. *, *p* < 0.05; n.s., not significant.

Next, we tested if these knockout cell lines could efficiently target Mtb to selective autophagy. We infected with mCherry Mtb, stained for the autophagy marker LC3, and quantified the percentage of targeted bacteria. Compared to control cell lines (GFP sgRNAs), galectin-8^-/-^ cell lines had less (approximately 50% less) LC3+ bacteria at 6 h post-infection (Fig. 3B, top). This was specific to galectin-8 as galectin-3^-/-^ and galectin-9^-/-^ cell lines had similar percentages of LC3+ Mtb compared to controls. These cell lines all had similar proportions of ubiquitin+ Mtb (Fig. 3B, bottom), which suggests that galectin recruitment is independent of ubiquitination.

To test how this defect in targeting impacts Mtb survival/replication in macrophages, we measured bacterial replication using an Mtb strain constitutively expressing luxBCADE. With this strain, at various timepoints post-infection, we could use luminescence as a proxy to monitor Mtb replication in numerous cell lines (Budzik et al., 2020; Hoffpauir et al., 2020; Penn et al., 2018). In control cells, Mtb replication is well-controlled; after 24 h bacterial burdens decrease before Mtb begins to slowly replicate intracellularly at later time points (Fig. 3C). However, in galectin-8^-/-^ macrophages, but not galectin-3^-/-^ or galectin-9^-/-^ macrophages, Mtb was not controlled at 24 h post-infection and instead replicated ~2-fold (Fig. 3C). In addition, the higher bacterial burdens in galectin-8^-/-^ cells persisted over 72 h of infection. This indicates that the defective selective autophagy targeting in galectin-8^-/-^ macrophages results in diminished control of Mtb survival/replication, and together, these data suggest that galectin-8 in particular is required for targeting Mtb to antibacterial selective autophagy.

Because several galectins are recruited to Mtb during infection, we next investigated whether they served redundant functions in targeting Mtb to selective autophagy. We used a lentiviral sgRNA array construct to simultaneously express the most efficient galectin-specific sgRNAs or as a negative control, GFP sgRNAs, in FLAG-Cas9-expressing RAW 264.7 cells. As with the single knockout lines, we isolated clonal cell populations, confirmed homozygous mutation of all three galectin genes, and validated the triple knockout cell lines by measuring galectin transcript levels (Fig. S1D). We infected the galectin-3/8/9^-/-^ triple knockout cells and GFP sgRNA control cells with mCherry Mtb and used immunofluorescence microscopy to quantify selective autophagy targeting. Compared to controls, the galectin-3/8/9^-/-^ cell lines had fewer LC3+ Mtb 6 h post-infection, but similar numbers of ubiquitin+ Mtb (Fig. 3D). When infected with luxBCADE Mtb, the galectin-3/8/9^-/-^ cell lines also had higher Mtb survival/replication compared to controls (Fig. 3E-F). Surprisingly, the magnitude of the defect in the galectin-3/8/9^-/-^ triple knockout cells (~2-fold defect) phenocopied that of the galectin-8^-/-^ single knockout lines (~2- to 2.5-fold defect), suggesting that these three galectins do not serve redundant functions, and instead galectin-8 has a privileged role in targeting Mtb to selective autophagy.

### Galectin-8 interacts with diverse proteins involved in exosome secretion, membrane trafficking, and selective autophagy

To gain a deeper understanding of how galectin-8 promotes targeting of Mtb to selective autophagy, we used an unbiased mass-spec approach. We predicted that galectin-8 may have one or more specific binding partners that would help explain why loss of galectin-8 in particular decreased LC3 recruitment to the Mtb-containing phagosome. Due to technical limitations resulting from Mtb’s classification as a Biosafety Level 3 (BSL3) pathogen, we turned to *Listeria monocytogenes*, a BSL2 pathogen that also elicits a type I IFN response, can be targeted to selective autophagy, and recruits galectins-3, -8, and -9 (Manzanillo et al., 2012; Mitchell et al., 2015; Thurston et al., 2012). To increase the population of *L. monocytogenes* targeted to selective autophagy, we used a strain lacking ActA, a protein that enables mobility within the host cell and therefore helps bacteria evade autophagy (Mitchell et al., 2015). We infected RAW 264.7 cells stably expressing 3xFLAG-galectin-8 with ΔactA *L. monocytogenes* at a multiplicity of infection (MOI) of 5, and immunoprecipitated 3xFLAG-galectin-8. Proteins associated with galectin-8 were identified using LC/MS (Fig. 4A).

**Figure 4.**
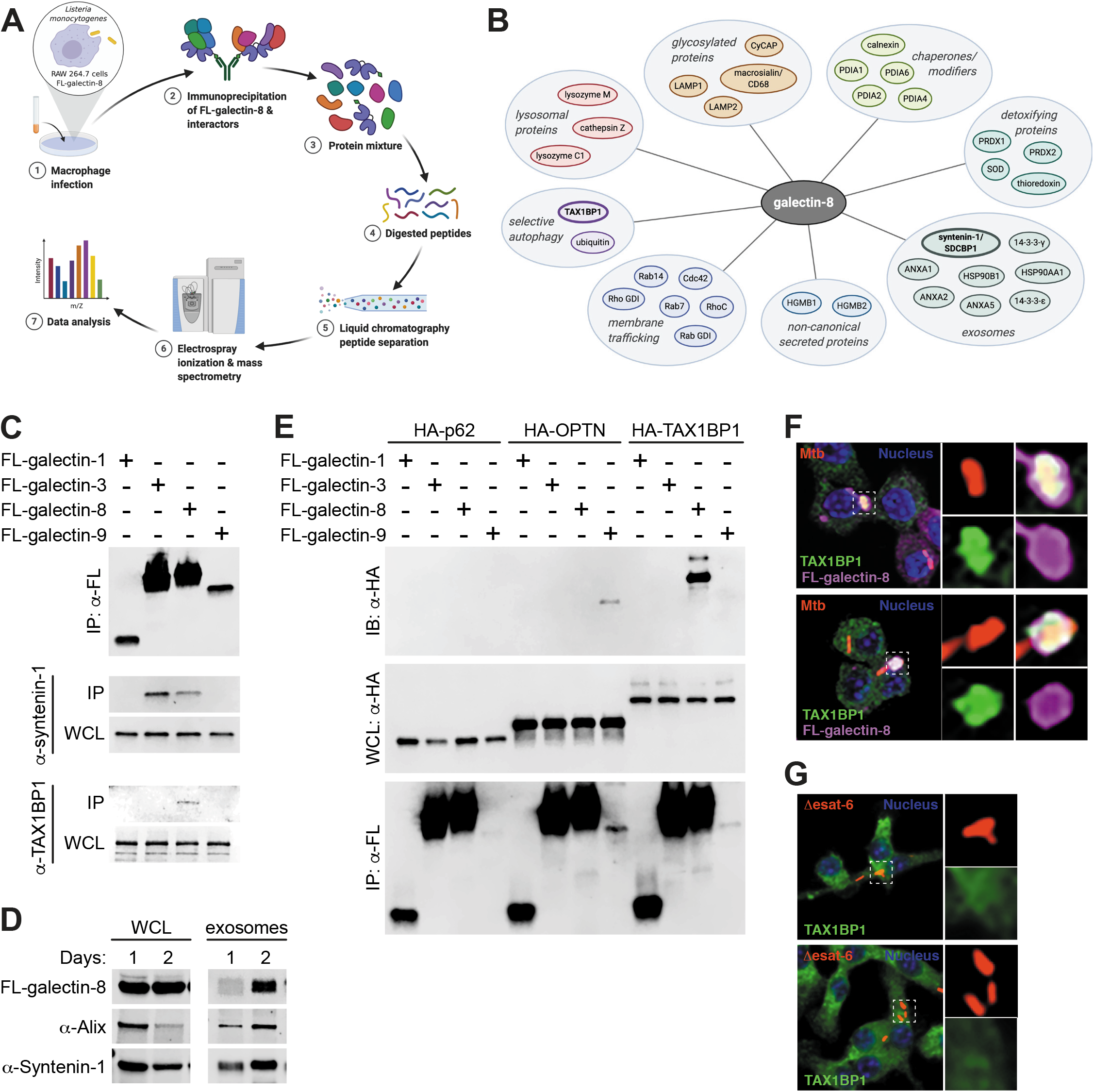
Galectin-8 interacts with exosome-associated proteins and selective autophagy adapter TAX1BP1. **(A)** Schematic of approach for immunoprecipitation and mass spectrometry (IP-MS) identification of galectin-8 binding partners in macrophages during in-tracellular bacterial infection. **(B)** Proteins identified by IP-LC/MS with galectin-8. **(C)** Co-immunoprecipitation (IP) of 3xFLAG (FL)-tagged galectins ectopically expressed in HEK293T cells. Whole cell lysates (WCL) and co-IPs probed for endogenous syntenin-1 and TAX1BP1. **(D)** WCL and exosomes from FL-galectin-8-expressing RAW 264.7 cells cultured for indicated number of days to assess exosome accumulation in cell culture media. **(E)** Directed co-IPs of FL-galectins and HA-tagged selective autophagy adapters. **(F)** Immunofluorescence of RAW 264.7 cells stably expressing FL-galectin-8 and co-stained for endogenous TAX1BP1 infected with WT mCherry-expressing Mtb (MOI = 1) 6 h post-infection. Green, endogenous TAX1BP1; magenta, FL-galectin-8; red, mCherry Mtb; blue, DAPI. **(G)** Immunofluorescence of RAW 264.7 cells infected with Δesat-6 mCherry Mtb and stained for endogenous TAX1BP1 at 6 h post-infection. Green, endogenous TAX1BP1; red, Δesat-6 Mtb; blue, DAPI. Panels (a-B) made with BioRender.com.

The protein interacting partners identified by IP-LC/MS provided insight into several novel aspects of galectin-8 biology. First, consistent with galectin-8 recognizing damaged phagosomes, endosomes, and lysosomes, we found lysosomal proteins (cathepsin Z, lysozyme M, lysozyme C1), highly glycosylated proteins (LAMP1, LAMP2, macrosialin/CD68, cyclophilin C-associated protein), chaperones/modifiers of glycosylated proteins (calnexin, protein disulfide isomerases [PDIA1, PDIA3, PDIA4, PDIA6]), and detoxifying enzymes (thioredoxin, superoxide dismutase, peroxireductases [PRDX1, PRDX2]). Additionally, we identified several galectin-8 binding partners with known roles in membrane trafficking (Rab7, Rab14, RhoC, Cdc42, Rab GDI [GDP dissociation inhibitor], Rho GDI) and cytoskeleton rearrangements (EFhd2, profilin, talin-1, gelsolin, F-actin capping proteins, macrophage capping protein), which are all consistent with galectin-8’s role in recognizing damaged endosomes and lysosomes.

Interestingly, we identified several proteins that, like galectins, are secreted through a non-canonical pathway that does not require a signal sequence, including HGMB1 (Gardella et al., 2002; Li et al., 2020). Also identified were a panel of proteins associated with exosome secretion, a form of non-canonical secretion, including syntenin-1/SDCBP, HSP90AA1, HSP90B1, ANXA1, ANXA2, ANXA5, 14-3-3-epsilon/YWAHAE, and 14-3-3-gamma/YWAHAG (Baietti et al., 2012; Gonzalez-Begne et al., 2009; Guha et al., 2019; Lauwers et al., 2018). Using co-immunoprecipitations of 3xFLAG-tagged galectins ectopically expressed in HEK293T cells, we confirmed this interaction between galectin-8 and endogenous syntenin-1 (Fig. 4C). Interestingly, this interaction was not unique to galectin-8 since galectin-3, but not galectins-1 or -9, also interacted with syntenin-1. These observations led us to hypothesize that galectin-8 could be secreted via exosomes. To test this, we isolated exosomes from the cell culture supernatant of RAW 264.7 cells and found that 3xFLAG-galectin-8, along with the exosomal proteins Alix and syntenin-1, were present in exosome preps (Fig. 4D). Moreover, the amount of exosomal galectin-8, Alix, and syntenin increased over time as exosomes accumulated in the cell culture media. Together, these data suggest that release in exosomes may be a key mechanism of secretion for extracellular galectins.

Finally, our mass spectrometry analysis identified ubiquitin, which is consistent with our observation that galectin-8 colocalizes with ubiquitin+ Mtb (Fig. 2), and it corroborates recent studies using global proteomics approaches that found galectin-8 itself is ubiquitinated during Mtb infection (Budzik et al., 2020; Penn et al., 2018). We also identified TAX1BP1 as a galectin-8 interacting protein. While TAXBP1 has been previously characterized as a selective autophagy adaptor with ubiquitin- and LC3-binding domains, it is not known to interact with galectins. We hypothesized that galectin-8 could augment selective autophagy of Mtb by binding to TAX1BP1 and promoting recruitment of downstream autophagy machinery.

### Galectin-8 interacts with TAX1BP1 independently of ubiquitination

We first confirmed the galectin-8/TAX1BP1 interaction using HEK293T cells ectopically expressing 3xFLAG-galectins and found that endogenous TAX1BP1 immunoprecipitated specifically with galectin-8 (Fig. 4C). To further probe the specificity of the galectin-8/TAX1BP1 interaction, we generated HA-tagged expression constructs for several selective autophagy adaptors, including TAX1BP1, p62, and optineurin/OPTN. We then tested the interaction between each galectin and adaptor by co-expressing pairs in HEK293T cells and performing directed co-IPs. Remarkably, we found that galectin-8 specifically interacted with TAX1BP1 and no other adaptors, and HA-TAX1BP1 interacted specifically with only with galectin-8 and no other galectins (Fig. 4E). We also detected an unexpected but seemingly specific interaction between galectin-9 and OPTN (Fig. 4E). These highly specific protein-protein interactions are surprising since there is a high degree of similarity between galectins (Fig. S2 A-B).

We next examined the localization of TAX1BP1 during Mtb infection. We infected RAW 264.7 cells expressing 3xFLAG-galectin-8 with mCherry Mtb and used immunofluorescence microscopy to visualize endogenous TAX1BP1. TAX1BP1 colocalized with galectin-8+ Mtb (Fig. 4F), and we found near complete overlap in the TAX1BP1+ and galectin-8+ populations. Furthermore, in cells infected with Δesat-6 Mtb, TAX1BP1 did not colocalize with Mtb, indicating that like galectins (Fig. 1C) and other adapters (Watson et al., 2012), phagosomal damage and/or cytosolic exposure is required for the recruitment of TAX1BP1 (Fig. 4G). Because we detected an interaction between galectin-9 and OPTN, we performed similar experiments co-staining for OPTN. However, we found that endogenous OPTN did not colocalize with Mtb at any time points examined (Fig. S2C). However, when we stably expressed 3xFLAG-OPTN in RAW 264.7 cells, we observed low levels of colocalization (Fig. S2C), suggesting that while OPTN is capable of being recruited to the Mtb-containing phagosome, it is unlikely to play a substantial role in the early targeting of Mtb to selective autophagy under normal conditions. Because several galectins (galectins-3, -8, and -9) and selective autophagy adapters (TAX1BP1, p62) are all recruited to the same population of Mtb-containing phagosomes, the highly specific galectin-8/TAX1BP1 interaction is particularly noteworthy.

To investigate the mechanisms by which these proteins interact, we made a series of truncations of both galectin-8 and TAX1BP1. TAX1BP1 contains several annotated domains, including a SKICH domain, an LC3-interacting region (LIR), a large coiled-coil domain, and two ubiquitin-binding zinc fingers (UBZs)(Fig. 5A). Because galectin-8 itself is likely ubiquitinated during infection, we predicted that TAX1BP1 binds galectin-8 via its UBZ domains. Surprisingly, when we performed directed co-IPs between galectin-8 and a panel of TAX1BP1 truncations, we found that the UBZ domains of TAX1BP1 were dispensable for its interaction with galectin-8 in this system (Fig. 5B). Instead, only the coiled-coil domain was required for interaction. To further narrow the region required for interaction with galectin-8, we tested additional truncations of TAX1BP1 that included combinations of the N- and C-terminals of the coiled-coil domain, an annotated oligomerization domain, and three smaller coiled-coil domains (Fig. 5A). In co-IPs with galectin-8 and these additional TAX1BP1 truncations, we found that the C-terminal portion of the coiled-coil domain was required and sufficient for this interaction (Fig. 5C). We propose calling this region of TAX1BP1 the galectin-8-binding domain (G8BD)(Fig. 5A). We next investigated truncations of galectin-8, which contains two carbohydrate recognition domains (CRDs) that are connected by a short flexible linker (Fig. 5D). In directed IPs, we found that the C-terminal CRD domain (CRD2), but not the N-terminal CRD (CRD1), interacted with TAX1BP1 (Fig. 4E). Together, these biochemical experiments indicate that TAX1BP1 has evolved a ubiquitin-independent mechanism to specifically interact with galectin-8.

**Figure 5.**
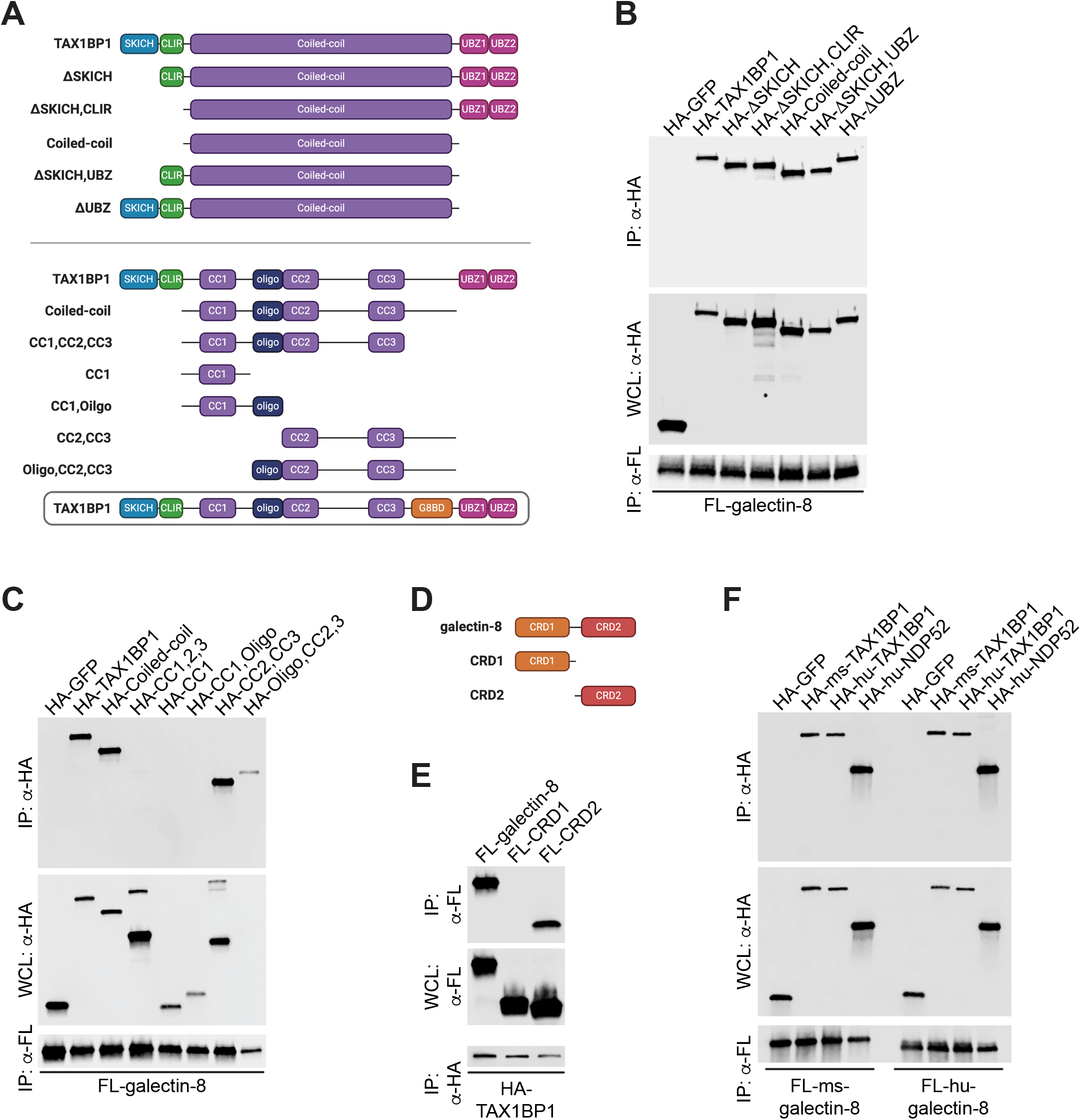
TAXIBP1’s coiled-coil domain and galectin-8’s CRD2 are required for their interaction. **(A)** Schematic representation of TAX1BP1 domain structure and truncations used in (B-C). CLIR, noncanonical/LC3C-interacting region; UBZ, ubiquitin-binding zinc finger domain; CC, coiled-coil domains; Oligo., oligomerization domain. **(B-C)** Directed co-immunoprecipitations (IP) of 3xFLAG (FL)-tagged galectin-8 ectopically expressed in HEK293Ts. Whole cell lysates (WCL) and co-IPs probed for HA-tagged TAX1BP1 truncations. HA-GFP shown as negative control for interaction. **(D)** Schematic of galectin-8 domain structure and truncations. CRD, carbohydrate recognition domain. **(E)** Directed co-IPs of HA-TAX1BP1 expressed in HEK293T cells. WCLs and co-IPs probed for FL-galectin-8 truncations. **(F)** As in (B-C) but with mouse (ms) and human (hu) FL-galectin-8, HA-TAX1BP1, and HA-NDP52. Panels (A) and (D) made with BioRender.com.

A previous study found that in non-immune cells, galectin-8 interacts with another selective autophagy adaptor, NDP52, which has a domain structure highly similar to TAX1BP1 (Fig. S2D)(Thurston et al., 2012). This study found that, similar to our findings in TAX1BP1, human NDP52 interacts with galectin-8 via the C-terminal region of NDP52’s comparatively smaller coiled-coil domain. Because of these similarities, we wanted to test the conservation of the TAX1BP1/galectin-8 interaction. To do this, we co-expressed human 3xFLAG-galectin-8 with human HA-TAX1BP1 or human HA-NDP52 and performed co-IPs. Consistent with previous reports, galectin-8 interacted with NDP52 (Fig. 5F). Importantly, human galectin-8 also interacted with human TAX1BP1 (Fig. 5F). This previously unidentified interaction indicates that galectin-8 can interact with both NDP52 and TAX1BP1 in humans. Based on our previous studies, the mouse gene encoding NDP52 appears to be disrupted by repetitive elements and lacks the regions previously shown to interact with galectin-8. Therefore, while the reported interaction between NDP52 and galectin-8 is likely not at play in mouse cells, it appears that human cells have evolved galectin-8 binding partners that may serve redundant functions. Finally, mouse galectin-8 can interact with human TAX1BP1 and human NDP52, and human galectin-8 can interact with mouse TAX1BP1 (Fig. 5F), which suggests that galectin-8, TAX1BP1, and their biochemical interactions are highly conserved.

### TAX1BP1 can be recruited to Mtb-containing phagosomes by binding galectin-8 or ubiquitinated proteins

To assess how the galectin-8/TAX1BP1 interaction influences targeting of Mtb to selective autophagy, we looked at the recruitment of TAX1BP1 to Mtb in galectin-8^-/-^ cells. The percentage of TAX1BP1+ Mtb in both galectin-8^-/-^ and galectin-3/8/9^-/-^ cell lines was lower compared to controls (Fig. 6A-B). This defect in recruitment was specific to TAX1BP1, though, since the number of p62+ Mtb was similar in knockout and control cells (Fig. 6A-B).

Because a sizeable population of Mtb were TAX1BP1+ even in the absence of galectin-8, we next investigated how specific domains of TAX1BP1 might contribute to its colocalization with Mtb. We predicted that because TAX1BP1 has UBZ domains, perhaps it could be recruited to Mtb in the absence of galectin-8 by binding to other ubiquitinated substrates surrounding the Mtb-containing phagosome. To test this, we stably expressed full-length HA-TAX1BP1 or HA-TAX1BP1 lacking the UBZ domains (HA-TAX1BP1ΔUBZ) in control and galectin-8^-/-^ cell lines. Then, at 6 h post-infection with mCherry Mtb, we performed immunofluorescence microscopy to quantify the number of HA-TAX1BP1+ bacteria (Fig. 6C). Consistent with experiments in Fig. 6A, which examined endogenous TAX1BP1, full-length HA-TAX1BP1 was recruited less efficiently in galectin-8^-/-^ cells (Fig. 6D), again indicating the galectin-8/TAX1BP1 interaction is required for TAX1BP1 recruitment. Furthermore, in control cells expressing HA-TAX1BP1ΔUBZ, even fewer Mtb were TAX1BP1+ (Fig. 6D), suggesting that TAX1BP1’s ability to bind ubiquitinated substrates is also required for its recruitment to Mtb. Finally, in support of our prediction, HA-TAX1BP1 ΔUBZ was recruited least efficiently in galectin-8^-/-^ cells (Fig. 6D), suggesting that both binding capabilities are involved in recruiting TAX1BP1 to Mtb. The residual recruitment of TAX1BP1 to Mtb in the absence of both galectin-8 and UBZ domains could be mediated by TAX1BP1’s LIR (LC3 interacting region) or other interactions or oligomerization with endogenous wild-type TAX1BP1. Together, these data demonstrate that TAX1BP1 can be recruited to damaged Mtb-containing phagosome by at least two independent mechanisms: binding to galectin-8 via its coiled-coil domain and binding to ubiquitinated substrates.

**Figure 6.**
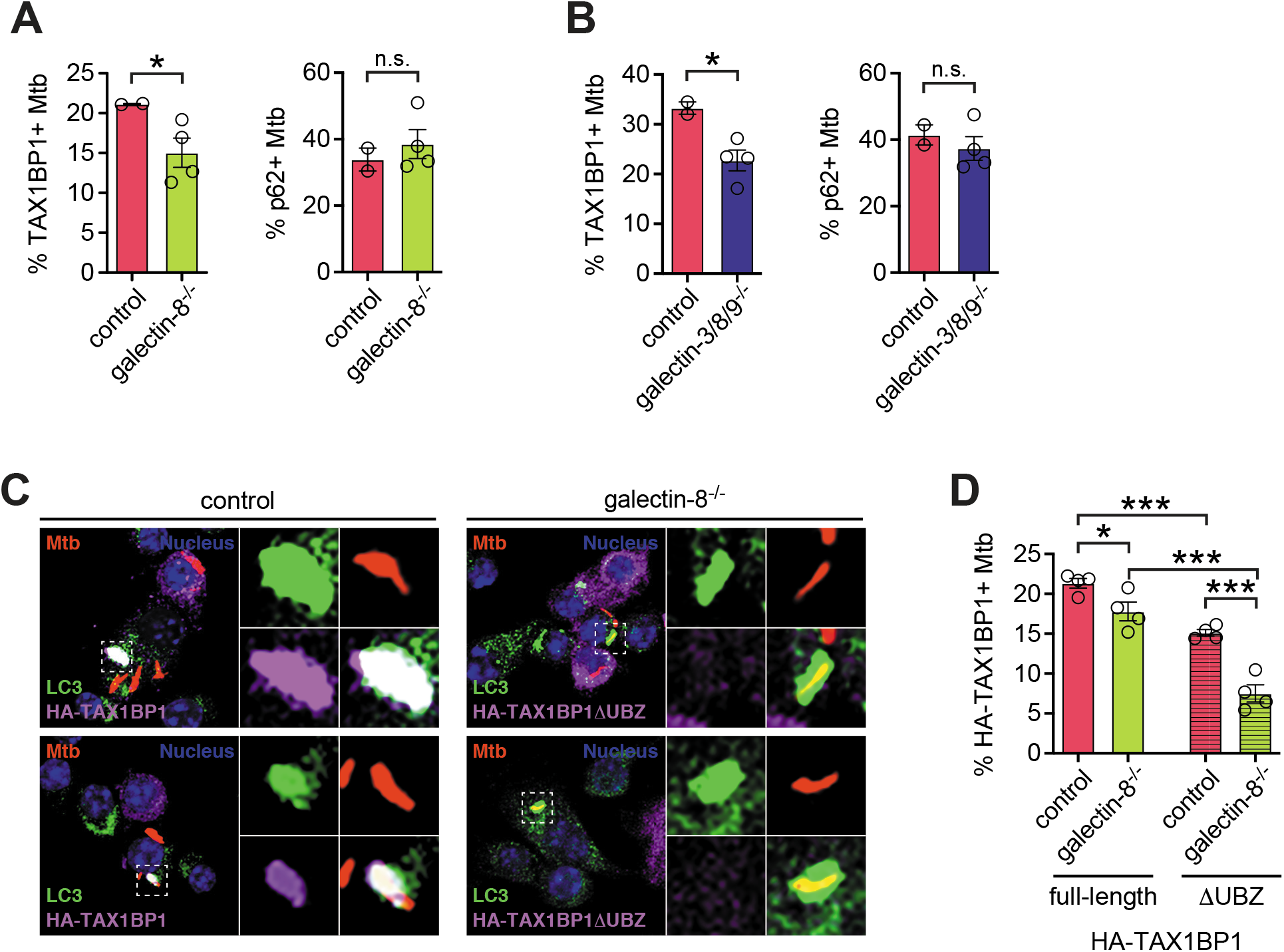
TAX1BP1 is recruited to Mtb-containing phagosomes by both its UBZ domain and its interaction with galectin-8. **(A)** Quantification of TAX1BP1- or p62-positive Mtb in control galectin-8 knockout RAW 264.7 cell lines at 6 h post-infection. Circles represent data for individual clonally selected cell lines. **(B)** As in (A) but with RAW 264.7 cell lines in which all three galectins are knocked out. **(C)** Immunofluorescence of control or galectin-8 knockout RAW 264.7 cells stably expressing full-length HA-TAX1BP1 or truncated HA-TAX1BP1 ΔUBZ that is missing its ubiquitin-binding domain. Cells were infected with WT mCherry-expressing Mtb (MOI = 1) and harvested at 6 h post-infection. Green, LC3; magenta, HA-TAX1BP1 variants; red, mCherry Mtb; blue, DAPI. **(D)** Quantification of indicated variant HA-TAX1BP1-positive Mtb in indicated genotype. Error bars indicate S.E.M. of knockout cell lines in which at least 300 bacteria per cell line were assessed. *, *p* < 0.05; *** *p* < 0.005; n.s., not significant.

### Overexpression of galectins augments targeting to selective autophagy

Finally, having characterized the requirement of galectins for targeting Mtb to selective autophagy, we next tested how overexpression of galectins might impact this pathway. RAW 264.7 cells overexpressing 3xFLAG-galectin-8 or galectin-9 had a small but significant increase in LC3+ Mtb at 6 h post-infection compared to cells overexpressing FL-galectin-1 (Fig. 7A). Importantly, the increased targeting in FL-galectin-8 and -9 cells translated to a significant increase in macrophages’ ability to control Mtb replication as measured by luxBCADE Mtb (Fig. 7B). Overexpression of FL-galectin-3 had a moderate effect on selective autophagy targeting and controlling Mtb replication/survival, which is consistent with its intermediate recruitment phenotype (Fig. 1A-B). Together, these data indicate that overexpression of galectins substantially enhances macrophages’ ability to recognize and respond to Mtb infection.

**Figure 7.**
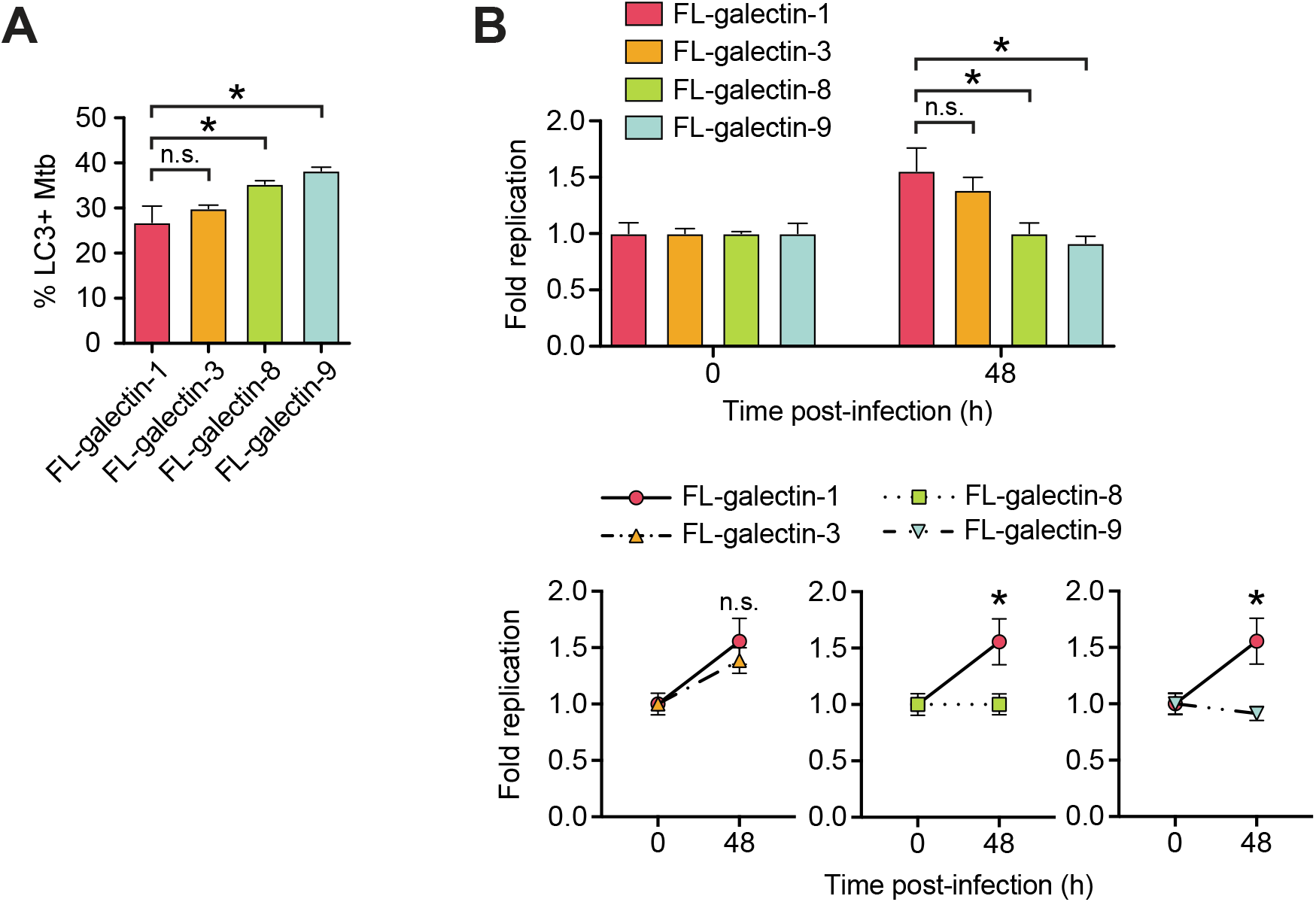
Overexpression of galectin-8 increases targeting and controls replication of Mtb. **(A)** Quantification of LC3-, TAX1BP1-, p62-, and ubiquitin-positive Mtb in RAW 264.7 cells overexpressing 3xFLAG (FL)-tagged galectins at 6 h post-infection. **(B)** Fold replication of luxBCADE Mtb (MOI = 1) in FL-galectin overexpression cell lines at indicated time points. Data normalized to t=0 h. Error bars indicate S.D. of overexpression cell lines; for IF, at least 300 bacteria per cell line were assessed. *, *p* < 0.05; n.s., not significant.

## DISCUSSION

Selective autophagy is a critical pathway employed by macrophages to control Mtb infection. Here we characterized the involvement of galectins, a family of damage/danger sensors, in the selective autophagy response to Mtb (Fig. 8). Of the galectins we studied, we found that galectin-8, but not galectin-3 or -9, was required for controlling Mtb infection in macrophages. This is somewhat surprising since all three galectins were recruited to the phagosome. However, the specific requirement of galectin-8 may be due to its interaction with the selective autophagy adapter TAX1BP1, which a recent report found to be required for targeting Mtb to selective autophagy and controlling Mtb replication in macrophages (Budzik et al., 2020). Our data indicate that TAX1BP1 can be recruited to the Mtb-containing phagosome in two ways: by binding directly to galectin-8, which is recruited directly to damaged Mtb-containing phagosomes, and by binding to ubiquitinated substrates. Consistently, this two-pronged recruitment of an adaptor, via galectin-8 and via ubiquitinated substrates, has been observed for NDP52 in HeLa cells infected with *S*. Typhimurium (Thurston et al., 2012). Since NDP52 and TAX1BP1 are highly related selective autophagy adapter it is perhaps not surprising that they have similar functional profiles. The curious similarities and apparent redundancies between adapters emphasize the importance of understanding the nature of their specific biological functions. Many important questions remain to be explored, including whether TAX1BP1 and NDP52 serve truly redundant roles in human autophagy or if one has evolved a particularly important function in other cell types.

**Figure 8.**
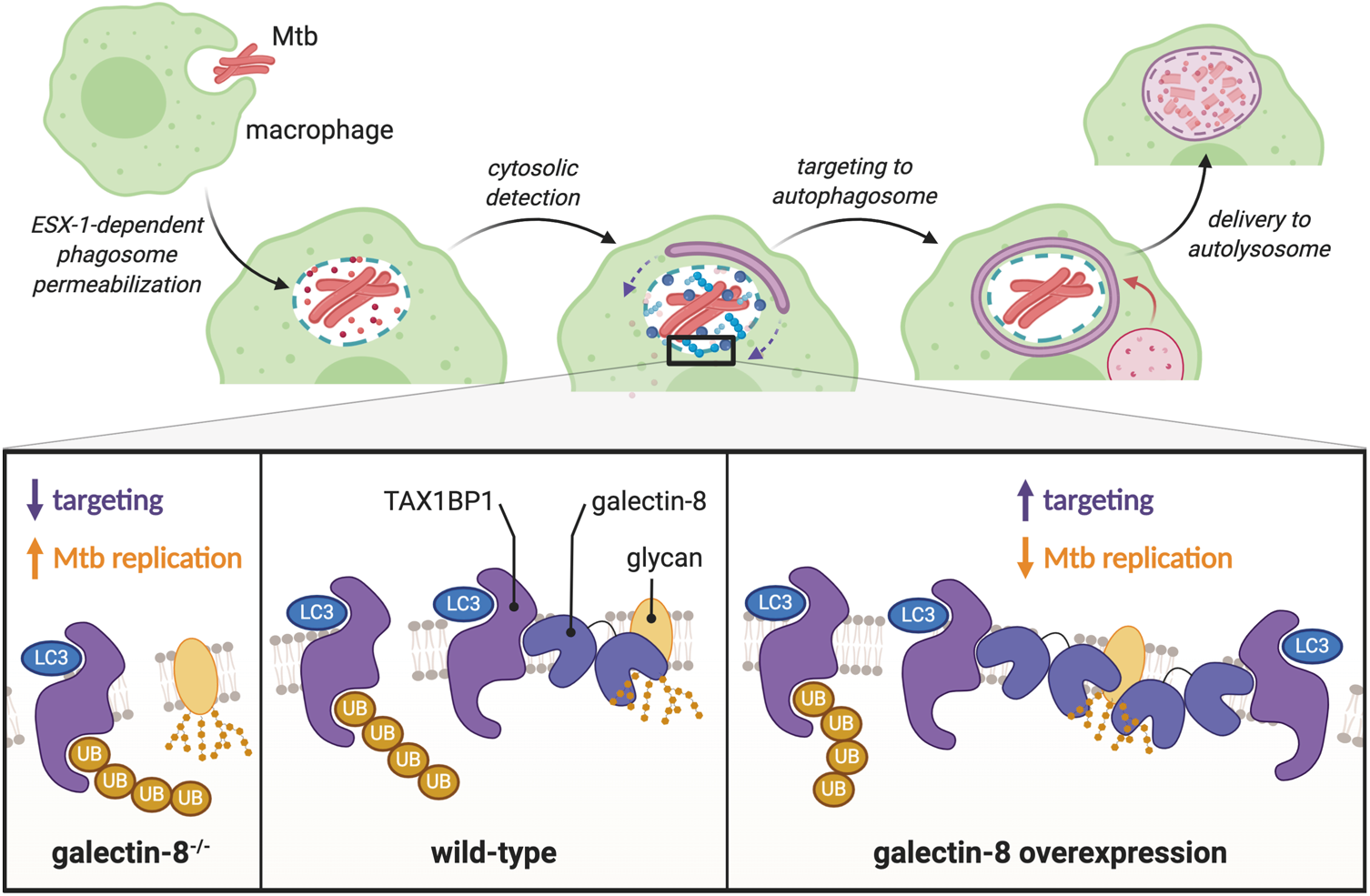
Galectin-8 and TAX1BP1 recognize and target Mtb to selective autophagy in macrophages. Schematic representation of how Mtb is detected by macrophages. Galectin-8 binds to cytosolically exposed glycans in the lumen of damaged Mtb-containing phagosomes. TAX1BP1 is recruited to these damaged phagosomes via its interaction with galectin-8 as well as through its interaction with ubiquitinated substrates. Deletion of galectin-8 results in less targeting of Mtb and increased survival/replication, while overexpression of galectin-8 leads to increased targeting and less Mtb replication. Made with BioRender.com.

Our experiments demonstrate that even early during infection, when Mtb appears to be enclosed inside a vacuole, there is sufficient disruption of the phagosomal membrane to permit entry of host factors into the lumen of the Mtb-containing phagosome. As a result, there is likely substantial exposure of both pathogen-associated and damage-associated molecular patterns (PAMPs and DAMPs) very early during Mtb infection. Some of the host pattern recognition receptors that detect these danger signals are known, including cGAS and now galectins, and these studies indicate that the molecular environment around the Mtb-containing phagosomes is extremely complex and is crowded with many proteins involved in various host response pathways: cGAS (STING/TBK1/TRIM14/IRF3), galectins, and ubiquitin (adapters [p62, TAX1BP1, NDP52, NBR1], LC3s/GABARAPs, E3 ubiquitin ligases [Parkin, TRIM16, Smurf]). However, the molecular mechanistic links between these different proteins and pathways remain somewhat obscure. As a kinase, TBK1 can phosphorylate adapters like OPTN, NDP52, and p62 (Richter et al., 2016; Wild et al., 2011), and phosphorylation of OPTN by TBK1 can increase its affinity for ubiquitin. However, it remains unknown whether TBK1 activation influences adapters’ affinity for ubiquitin, LC3, or galectins during Mtb infection. Furthermore, the E3 ubiquitin ligases Parkin, Smurf1, and TRIM16 colocalize with Mtb and contribute to the ubiquitin cloud surrounding bacteria, but how these E3 ligases are activated upon Mtb infection and what proteins each E3 modifies are lingering unanswered questions. Finally, since Mtb is an exquisitely evolved pathogen, it is very likely that yet-to-be-identified bacterial proteins are intimately involved in these processes. Indeed, the recent discovery of a ubiquitin-binding protein (Rv1468c) on Mtb’s surface suggests that Mtb does have mechanisms for modulating the host’s selective autophagy pathway (Chai et al., 2019).

In our studies, we found that galectin-8 is required for targeting Mtb to selective autophagy. However, removal of this danger sensor did not completely abrogate targeting. This parallels what we have seen when the DNA sensor cGAS is depleted; around 50% of Mtb bacilli are still targeted (Watson et al., 2015). There are several possible explanations for this. First, these two pathways may function in parallel, each targeting some fraction of Mtb bacilli, adding up to the total of ~30% Mtb targeted in a wild-type cell. Future studies in cells lacking both cGAS/STING and galectin-8 could address this possibility. Second, it is possible that Rv1468c, Mtb’s ubiquitin-binding surface protein, contributes substantially to the ubiquitin cloud, and because of Rv1468c, removing host sensors will only ever decrease targeting to ~15% (Chai et al., 2019). It is likely that some of the Rv1468c-bound ubiquitin chains serve as substrates to recruit TAX1BP1 and other adapters, so using Rv1468c mutants in future studies of host sensing pathways will help elucidate if additional host factors remain to be discovered in the targeting of Mtb. Ultimately, understanding the molecular mechanisms underpinning the host-pathogen interactions between this sort of Mtb protein and macrophage proteins will be critical for understanding the innate immune response to Mtb.

Previous studies of galectins have examined the in vivo requirement for individual galectins during Mtb infection. Interestingly, they have found that galectin-8^-/-^ and galectin-3^-/-^ mice succumb more rapidly to Mtb infection, suggesting these galectins are required for controlling Mtb infection (Chauhan et al., 2016, p. 3; Jia et al., 2018). However, these studies did not further interrogate how galectins contributed to innate immunity during Mtb infection, and galectins are multifunctional proteins that play a multitude of roles *in vivo* beyond their intracellular function in macrophages. To really understand how individual galectins contribute to macrophages’ ability to control Mtb *in vivo*, future studies will need to infect mice with macrophage-specific deletions of these galectins. Such experiments would provide some of the best evidence to date of how selective autophagy in particular (rather than bulk autophagy) contributes to the control of Mtb infection *in vivo*. Of note, a previous study infected p62^-/-^ mice with Mtb but found no differences between wild-type and p62^-/-^ mice; however, as demonstrated here and in other recent studies, several selective autophagy adapters are involved in detecting and targeting Mtb, so it is likely that removing multiple adapters will be necessary in order to study the *in vivo* requirement of selective autophagy adapters.

Finally, the finding that overexpression of galectins can enhance macrophages’ ability to control Mtb is particularly noteworthy. Several host and bacterial factors can be mutated to diminish the targeting of Mtb to selective autophagy, but there are few known ways to enhance this targeting. In fact, for many other intracellular bacterial pathogens like Mtb, the targeted percentage is rarely above ~30%, suggesting this might be a biological setpoint that is difficult to overcome. However, it seems that galectin overexpression, even at the moderate levels permitted by our lentiviral expression system (Fig. S1B), is able to accomplish this. Identifying a class of proteins like galectins that can enhance targeting without causing significant off-target effects is extremely valuable in the future development of anti-TB therapies. For instance, overexpression or stimulation of cGAS, which is required for targeting, may enhance the number of targeted Mtb bacilli, but chronic activation of cGAS also results in enhanced production of type I IFNs, which are pro-bacterial and cause increased disease pathology *in vivo*. While chronic overexpression of galectins can have detrimental effects (Vinik et al., 2015), using small molecules to augment the function of galectins specifically during infection might be an especially attractive strategy for the future development of host-directed therapies for TB.

## MATERIALS AND METHODS

### Cell lines and cell culture

RAW 264.7 cells (ATCC TIB-71) and HEK293T cells (ATCC CRL-3216) were cultured in DMEM + 10% heat inactivated FBS + HEPES at 37 C with 5% CO2. Lenti-X (Takara Bio) cells were used for producing lentiviral particles. Where necessary, RAW 264.7 cells were selected with and maintained in Puromycin (InvivoGen, 5 μg/ml), Blasticidin (InvivoGen, 5 μg/ml), G418/Geneticin (InvivoGen, 750 μg/ml), or Hygromycin B (Life Sciences, 100 μg/ml). For infections, antibiotics were omitted from culture media.

RAW 264.7 cells were plated at 2x10^5^ cells/well in on circular glass coverslips in 24-well tissue culture (TC) dishes for immunofluorescence experiments and at 3x10^5^ cells/well in 12-well TC dishes for luciferase growth assays.

Epitope-tagged expression constructs were made by first cloning cDNAs from RAW 264.7 cell RNA into pENTR1a entry vectors with indicated tags (Addgene Plasmid #17396)(Campeau et al., 2009; Hoffpauir et al., 2020, p. 14; Watson et al., 2015). Constructs were fully Sanger sequenced (Eton Biosceinces, San Diego, CA) to verify the tagged proteins were complete, in-frame, and error-free. Constructs were then Gateway cloned with LR Clonase (Invitrogen) into pLenti destination vectors (Addgene Plasmid #19067)(Campeau et al., 2009). Expression of tagged proteins was confirmed by transfecting HEK293Ts with 1 μg of pDEST and harvesting cell lysates after 1-2 days of expression. Proteins were separated by SDS-PAGE and visualized by Western blot analysis using primary antibodies for FLAG (Clone M2, Sigma-Aldrich, F1804) and HA (Roche, 11867423001).

To make RAW 264.7 stable expression cells lines, Lenti-X 293T cells (Takara Bio) were co-transfected with pLenti plasmids and the packaging plasmids psPAX2 and pMD2G/VSV-G (Addgene Plasmids #12259-60) to produce lentiviral particles. RAW 264.7 cells were transduced with lentivirus for two consecutive days plus 1:1000 Lipofectamine 2000 (Invitrogen) and selected for 3-5 days with antibiotic. Expression of tagged proteins was confirmed by Western blot analysis with antibodies against indicated tag.

### Bacterial infections

Erdman was used as the parental Mtb strain for these studies (Stanley et al., 2007; Watson et al., 2012, 2015). The wild-type mCherry, Δesat-6 mCherry, and luxBCADE strains have been described previously (Budzik et al., 2020; Hoffpauir et al., 2020; Penn et al., 2018; Watson et al., 2012, 2015). Mtb cultures were grown in Middlebrook 7H9 (BD Biosciences) + 10% BBL Middlebrook OADC (Becton Dickinson) + 0.5% glycerol + 0.1% Tween-80 at 37°C in roller bottles. Strains were propagated with minimal passage.

Mtb infections were performed as previously described (Hoffpauir et al., 2020; Stanley et al., 2007; Watson et al., 2015). Briefly, cultures grown to 0.6-0.8 OD_600_ were spun at 500*g* for 5 min to remove large clumps and then spun again at 3000*g* for 5 min to pellet bacteria. After washing twice with PBS, bacteria were resuspended in PBS, sonicated briefly to disrupt clumps, and then spun once more at 500*g* for 5 min to remove remaining clumps. The OD_600_ of the bacterial suspension was used to calculate the volume needed for the desired multiplicity of infection (MOI) of 1 (1 OD = 3x10^8^ bacteria/ml). Bacteria were diluted in DMEM + 10% horse serum and added to cells. Infections were synchronized by spinning for 10 min at 1000*g*, and cells were washed twice with PBS and cultured in regular media. When experiments lasted for more than 24 h, cell culture media was replaced daily. For IF experiments, at the indicated time points, coverslips were transferred to 4% fresh paraformaldehyde in PBS, fixed for 20 min, and washed three times with PBS. For luciferase experiments, cells were washed twice with PBS, lysed in 0.5% Triton X-100, and transferred to a white luminescence plate (LumiTrac 96-well plates, Greiner Bio-One). Luminescence was measured using a Tecan Infinite 200 PRO. For 0 h time point, cells were lysed after PBS washes rather than being returned to cell culture media.

*Listeria monocytogenes* infections were also performed as previously described (Watson et al., 2015). RAW 264.7 cells were plated at 1x10^8^ cells per plate in 10 cm dishes. *Listeria monocytogenes* ΔactA (parental strain 10403, gift from Dan Portnoy) was grown in BHI (BD) at 30°C overnight without shaking. Culture was diluted 1:10 in BHI and grown for 3-4 h at 37°C without shaking until it reached an OD_600_ of ~0.6. Bacteria were washed twice with HBSS, and the OD of the resulting bacterial suspension was used to calculate the volume needed for an MOI of 5 (1 OD=1x10^8^ bacteria/ml). Bacteria were diluted in HBSS and added to cells. After incubating cells and bacteria for 30 min at 37°C, cells were washed twice with HBSS + 40 μg/ml gentamycin and then cultured in media + 10 μg/ml gentamycin until harvest.

### CRISPR/Cas9 Knockouts

RAW 264.7 cells stably expressing FL-Cas9 were generated by transducing RAW 264.7 cells with lentivirus containing LentiCas9-Blast (Addgene plasmid #52962)(Sanjana et al., 2014). These cells were selected with 5 μg/ml Blasticidin (Invivogen) for 3-5 days and then with 10 μg/ml Blasticidin for an additional 1-2 days. FL-Cas9 expression was confirmed via western blot analysis.

sgRNAs for each galectin gene were designed using the sgRNA Designer: CRISPRko website (https://portals.broadinstitute.org/gpp/public/analysis-tools/sgrna-design) website and synthesized by IDT (Doench et al., 2016; Sanson et al., 2018). sgRNAs used for each galectin were as follows: gfp-1: gggcgaggagctgttcaccg; gfp-2: cagggtcagcttgccgtagg; gal3-1: tctggaaacccaaaccctca; gal3-2: ggctggttcccccatgcacc; gal8-1: tcagtaatggtgccaacata; gal8-2: cagtaatggtgccaacatag; gal9-1: taccctccttcctcaaaccg; gal9-2: acccccggtttgaggaagga. Primers were cloned into LentiGuide-Puro (Addgene plasmid #52963) by phosphorylating, annealing, and ligating primers into digested vector (Sanjana et al., 2014; Shalem et al., 2014). sgRNA plasmids were validated by Sanger sequencing using the universal pLKO.1/hU6 promoter primer (Eton Biosciences, San Diego, CA). Lentivirus with sgRNAs were produced and used to transduce low passage FL-Cas9 RAW 264.7 cells. After selection with 5 μg/ml puromycin, the knockout efficiency was assessed at the population level. Using cells from the two most efficient sgRNAs, individual cells were serially diluted and plated into 96 well dishes to isolate clonal populations. When clones grew, populations were expanded, and each was assayed for mutations by amplifying a 500bp segment of genomic DNA around the mutation. These PCR fragments were Sanger sequenced using nested primers and compared to controls using TIDE analysis (https://tide.deskgen.com). Clones with homozygous nonsense mutations were further validated by measuring galectin RNA expression.

Triple knockout lines were made using a modified multiplexed lentiviral sgRNA system (Kabadi et al., 2014). The Cas9 in the lentiviral plasmid from this system was replaced with the puromycin resistance gene from a pDEST plasmid, which allowed for drug selection of a sgRNA array in RAW 264.7 cells already expressing FL-Cas9. sgRNAs for individual galectin genes were cloned into the sgRNA expression plasmids and assembled via Golden Gate assembly into the lentiviral backbone as previous published (Kabadi et al., 2014). RAW 264.7 cells expressing FL-Cas9 were transduced with lentivirus containing sgRNA arrays (GFP sgRNAs or galectin sgRNAs), and cells were selected, cloned, and screened as above.

### Immunofluorescence

Coverslips with fixed cells were blocked and permeabilized in 5% non-fat milk in PBS + 0.1% saponin for 30 min. Coverslips were then stained with primary antibody diluted in PBS with 5% milk and 0.1% saponin for 2-4 h. Primary antibodies used in this study were FLAG (Clone M2, Sigma-Aldrich, F1804; 1:1000), FLAG (Sigma-Aldrich, F7425; 1:1000), HA (Roche, 11867423001; 1:1000), LC3 (Invitrogen, L10382; 1:250), ubiquitin (Clone FK2, Millipore Sigma, 04-263; 1:500), p62 (Bethyl, A302-855A; 1:500), TAX1BP1 (A303-791A; 1:500), and OPTN (Bethyl, A301-829A; 1:500). Coverslips were washed three times in PBS and stained with secondary antibodies (Goat anti-Rabbit Alexa Fluor 488, Goat anti-Rat Alexa Fluor 647, and/or Goat anti-Mouse Alexa Fluor 647; Invitrogen, 1:1000) and DAPI (1:10,000) in PBS + 5% milk + 0.1% saponin for 1-2 h. Coverslips were then washed twice with PBS and twice with water and mounted using Prolong Gold Antifade Mountant (Thermo Fisher). Cells were imaged on an Olympus Fluoview FV3000 Confocal Laser Scanning microscope. Three coverslips per genotype were imaged, and at least 300 bacteria per coverslip were assessed and counted.

### Immunoprecipitations

HEK293T cells were plated at 5x10^7^ cells per plate in 6cm TC dishes. The following day, cells were transfected with 2-5 μg of indicated expression plasmids using PolyJet (SignaGen) according to manufacturer’s instructions. Typically, 1 μg of bait plasmid and 1-4 μg of prey plasmid were cotransfected. After two days, cells were washed with PBS, lifted using PBS + EDTA, and pelleted by centrifuging at 3000*g* for 5 min. Cells were lysed in lysis buffer (150 mM Tris pH 7.5, 50 mM NaCl, 1 mM EDTA, 0.075% NP-40, protease inhibitors), and lysates were cleared of cellular debris and nuclei by spinning at 7000g for 10 min. 5% of the cleared lysate was saved as the “whole cell lysate”, mixed with 4x Laemmli sample buffer with fresh ß-mercaptoethanol (Bio-Rad), and boiled for 5 min. Remaining cell lysate was incubated with pre-washed (three times in 1 ml lysis buffer) 20 μl of antibody-conjugated beads/resin (FLAG: EZview Red ANTI-FLAG M2 Affinity Gel, Sigma-Aldrich; HA: Pierce Anti-HA Agarose, Thermo Scientific) for 30-60 in at 4°C with rotation. Beads were washed three times with 1 ml wash buffer (150 mM Tris pH 7.5, 50 mM NaCl, 1 mM EDTA, 0.05% NP-40), and proteins were eluted with an excess of FLAG peptide (Sigma) or HA peptide (Sigma) resuspended in lysis buffer + 1% NP-40. Eluates were mixed with 4x sample buffer and boiled for 5 min. Proteins in whole cell lysates and immunoprecipitations were resolved by SDS-PAGE and imaged by western blot analysis using FLAG or HA antibodies (1:5000 in Li-Cor TBS Blocking Buffer), corresponding Li-Cor secondary antibodies (1:15,000), and a Li-Cor Odyssey Fc imager. Immunoprecipitations in RAW 264.7 cells stably expressing 3xFLAG-tagged proteins were performed using the same protocol and workflow using cells infected with *Listeria monocytogenes* 1 or 2 h post-infection.

### RNA extraction and RT-qPCR

Cells were harvested in Trizol and RNA was extracted using Direct-Zol RNA MiniPrep kits (Zymo Research) with at least 1 h of on-column DNase treatment. cDNA was made using iScrpit (Bio-Rad), and gene expression was quantified using relative standard curves on a QuantStudio 6 Flex (Applied Biosystems) with PowerUp SYBR Green Master Mix (Applied Biosystems). Primers for *Actb* (F-ggtgtgatggtgggaatgg, R-gccctcgtcacccacatagga), *Lgals3* (F-ctggaaacccaaaccctcaa, R-aggagcttgtcctgggtag), *Lgals8* (F-ccctatgttggcaccattact, R-gctgaaagtcaacctggaatct), and *Lgals9* (F-gcccagtctccatacattaacc, R-gttctgaaagttcaccacaaacc) were synthesized by IDT.

### Exosomes

RAW 264.7 cells were plated at 5x10^7^ cells per plate in 10 cm dishes. After 1 or 2 days in culture, cell culture media was collected, and cells were washed once with PBS and harvested by scraping. For whole cell lysates (WCLs), cells were pelleted and lysed directly in 1x Laemmli sample buffer with fresh ß-mercaptoethanol (Bio-Rad), sonicated to break up DNA, and boiled for 5 min. Culture media was precleared of dead cells and cell debris by spinning for 5 min at 3000g. Exosomes were then collected by ultracentrifugation for 1 h at 100,000*g*. Exosome pellets were resuspended directly in 1x sample buffer and boiled for 5 min. Proteins from WCLs and exosomes were resolved and imaged by SDS-PAGE and western blot analysis as described above using antibodies for FLAG (Sigma, F-1804; 1:5000), Alix (Abcam, ab117600; 1:2500), and syntenin-1 (Abcam, ab19903; 1:2500).

### Data analysis and presentation

Statistical analysis was performed using Prism (GraphPad) with Student’s unpaired two-tail t-tests. Graphs were generated with Prism, figures with Adobe Illustrator and Photoshop, and diagrams and schematics with BioRender.com as indicated in figure legends. At least three independent experiments were performed, and data presented is representative of these experiments. Figure legends indicate whether error bars indicate standard deviation (S.D.) or standard error of the mean (S.E.M.).

## ACKNOWLEDEMENTS

We’d like to thank Dr. Larry Dangott at the Texas A&M Protein Chemistry Lab for his assistance with mass spectrometry analysis and Dr. Malea Murphy at the Texas A&M Health Science Center Integrated Microscopy and Imaging Lab for her help with microscopy. Kayla Lopez and Kendall Green assisted in preparing necessary reagents. Schematics and diagrams were made using BioRender.com, and additional design consultation was provided by Maryann M. Bell and Kelsi West. We thank the members of the Patrick/Watson Lab, as well Dr. James Samuel, Dr. Phillip West, and their respective labs for their helpful discussions and feedback.

## AUTHOR CONTRIBUTIONS

Conceptualization, R.O.W., S.L.B., and K.L.P.; Investigation, S.L.B., R.O.W., K.L.L.; Writing, S.L.B., K.L.P., and R.O.W.; Visualization, S.L.B. and R.O.W.; Funding acquisition, R.O.W., K.L.P., and J.S.C.; Supervision, S.L.B., R.O.W., K.L.P., and J.S.C.

---

**Figure S1.**
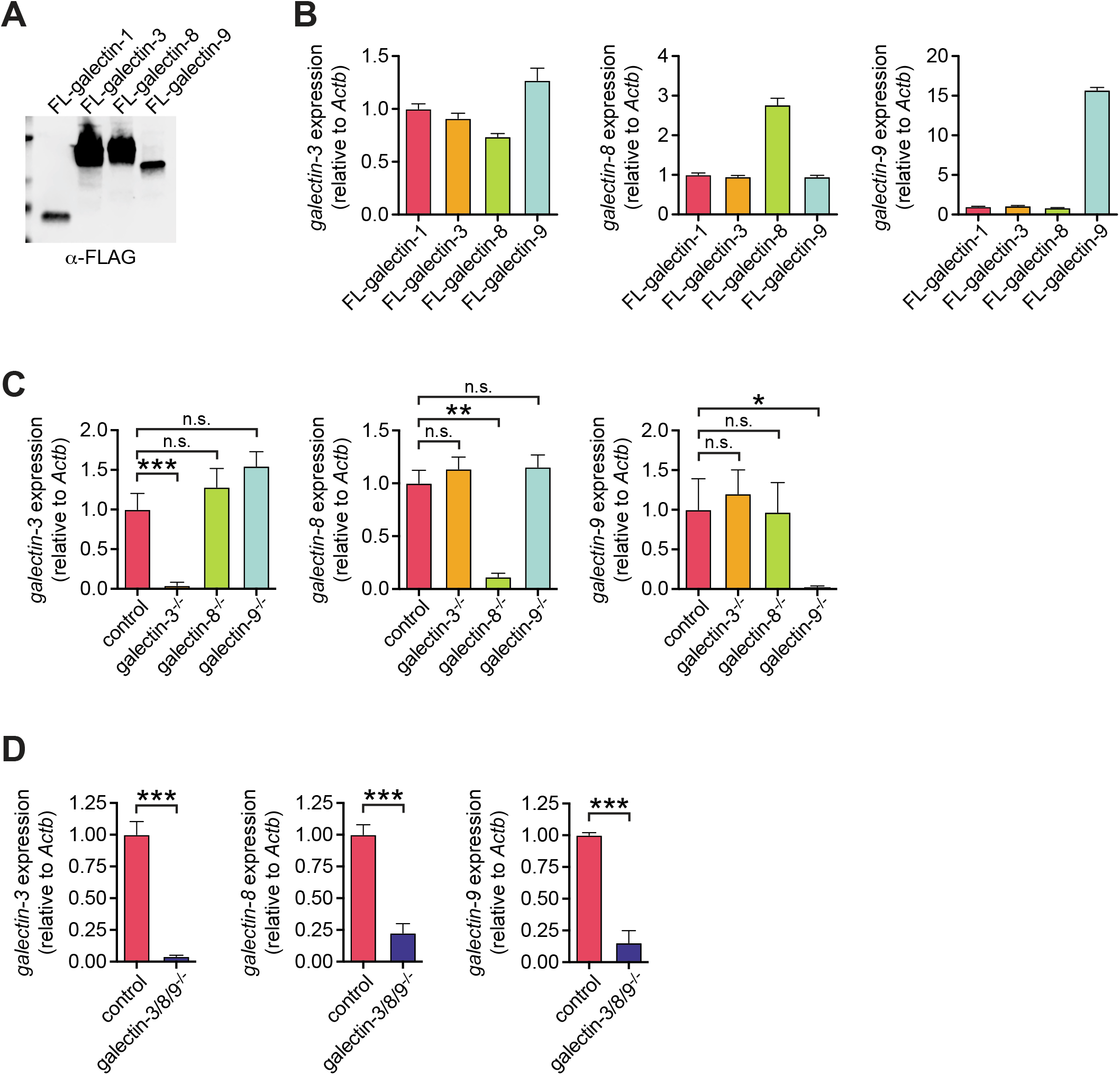
Validation of galectin stable expression and knockout cell lines. **(A)** Western blot of whole cell lysates from RAW 264.7 cells stably expressing indicated 3xFLAG-tagged galectins. **(B)** Transcript levels of indicated galectins in cells stably expressing epitope-tagged galectins in RAW 264.7 cells. **(C-D)** As in (B) but for control (GFP gRNA) and individual knockout cell lines (C) or control (GFP gRNA) and triple knockout cell lines (D). Error bars are S.E.M. of the averages of each of 3-5 clonal cell lines of each genotype. *, *p* < 0.05; ** *p* < 0.01; ***, *p* < 0.005; n.s., not significant.

**Figure S2.**
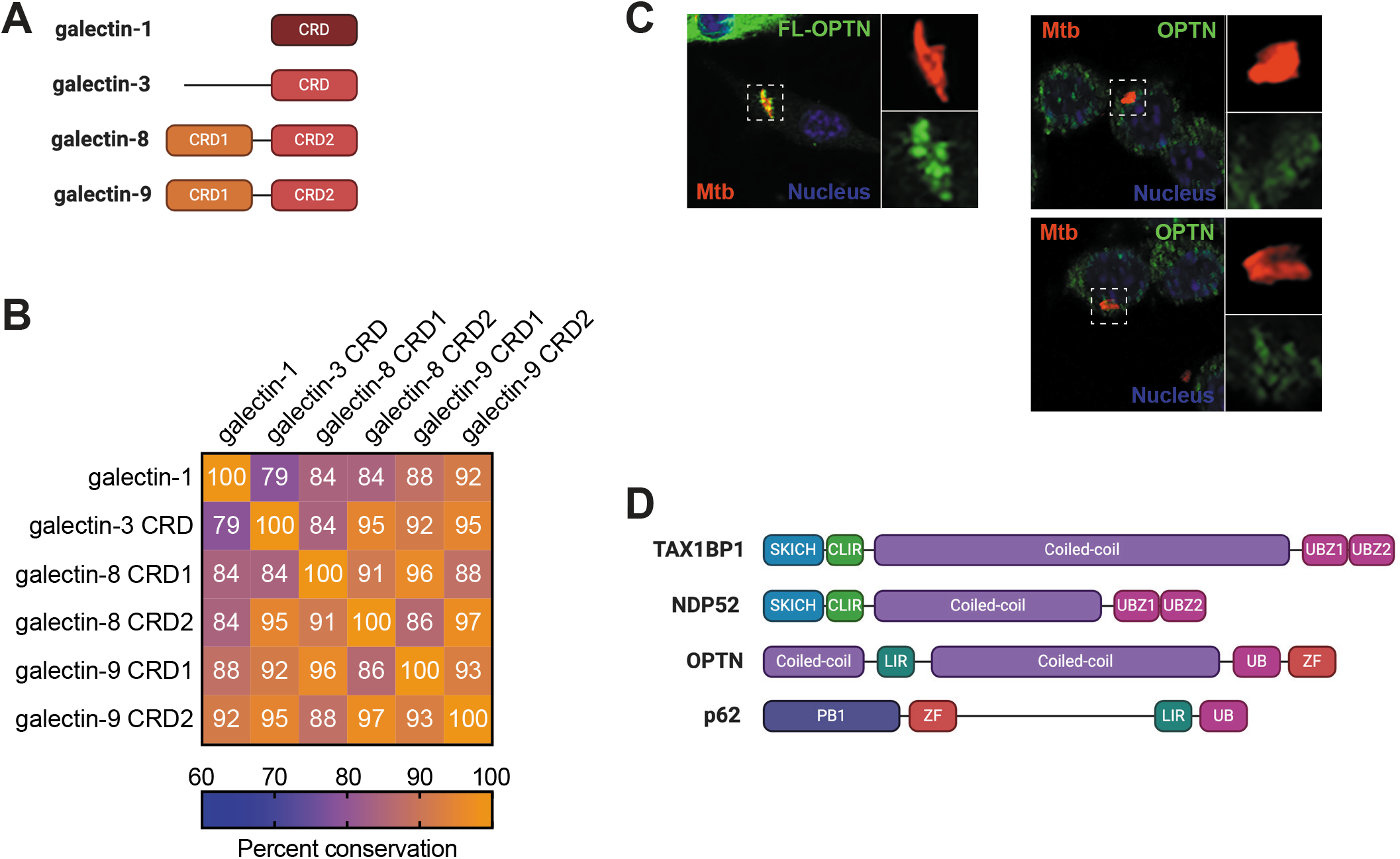
Domains of galectins and adapter proteins. **(A)** Schematic of the domains of galectin-1, -3, -8, and -9. CRD, carbohydrate recognition domain. **(B)** Percent conservation in pairwise comparisons of CRDs by analysis using M-Coffee. **(C)** Immunofluorescence of 3xFLAG-tagged OPTN (left) and endogenous OPTN (right) in RAW 264.7 cells infected with mCherry Mtb at 6 h post-infection. **(D)** Domain structure of selective autophagy adapters in this study. SKICH, coiled-coil, and PB1 are protein-protein interaction domains. CLIR, noncanonical/LC3C-interacting region; LIR, LC3-interacting region; ZF, zinc finger; UB, ubiquitin-recognition domain; UBZ, ubiquitin-binding zinc finger domain. Panels (A) and (D) made with BioRender.com.

